# Do light eaters live shorter lives? The case of ultralight *Caenorhabditis elegans*

**DOI:** 10.1101/2024.02.13.580069

**Authors:** Xuepei Zhang, Hassan Gharibi, Christian M. Beusch, Zhaowei Meng, Amir A. Saei, Massimiliano Gaetani, Roman A. Zubarev

## Abstract

The idea that ingesting heavy stable isotopes can increase longevity emerged shortly after the discovery of deuterium in the early 1930s and has been extensively tested since then on animals. Here we present the first experimental evidence for the opposite. Growing *C. elegans* on bacteria *E. coli* that are in turn fed on a diet depleted of heavy isotopes of C, H, N and O produced ultralight worms that grow and mature faster but have a shorter lifespan. Based on the differences in expression and solubility of proteins, we established an aging pseudo-time scale. Notably, the newly born ultralight worms appear to be significantly “younger” than their normal counterparts, while at day 10 they are significantly “older”. Pathway analysis revealed involvement of mitochondria; analysis of reactive oxygen species (ROS) confirmed significant ROS overproduction in ultralight worms that increases further with age. These findings provide a new modality of affecting the lifespan in this important animal model of human diseases and aging.

---

*Caenorhabditis elegans* (*C*.*elegans*) is a nematode feeding on bacteria that can be easily cultivated on agar plates or in liquid medium supporting *Escherichia coli*. Mature adults of this 1 mm long multicellular organism consist of 959 somatic cells for the hermaphrodites (99.8%) and 1031 cells for the males (0.2%), with the anatomical arrangement and entire cell lineage of all somatic cells being known for decades ^1^. The nematode life cycle lasts from egg through four larval stages ∼3.5 days at 20°C, after which the egg-laying adult lives up to 2–3 weeks under favorable conditions ^2^. *C. elegans* lends itself to non-invasive manipulation and optical monitoring, which helped in the investigation of the molecular mechanisms during the worm’s development. Since an estimated 60–80% of genes of *C. elegans* have homologs in humans, *C. elegans* is also being used as a model of human diseases and aging, including age-related neurodegenerative disorders^3^.

Modulation in research of the activity of metabolic and signaling pathways affecting nematode’s phenotype is achieved either via genetic engineering that alters the animal’s genome or transcriptome (e.g., RNAi-mediated knockdown ^4^), or by changing the growth environment, in particular diet ^5^. In studying the aging processes, the environment modulation approach is preferred, as it is less drastic and easier to interpolate to higher species ^6-8^.

The finding that dietary restriction (DR) can increase the lifespan of model organisms, including *C. elegans*^*9*^, has been met with a burst of enthusiasm ^10-12^. It has been found that different DR strategies extend nematode lifespan to different degrees independently of the calorie content ^11^. For instance, DR by bacterial food deprivation significantly increased life span in worms even when initiated at 24 days of adulthood, when >50% of the cohort have already died ^13^. Remarkably, it has been found that not only bacterial food itself, but its diffusible components (“smell of food”) affect worms’ life span ^14^.

The latter result underscores the complexity of studying such an apparently simple and well-understood system as *C. elegans* and calls for employing more subtle tools of its manipulation. One such tool is stable isotopes, as their biological effect, such as toxicity (except for deuterium at high concentration), is believed to be minimal ^15^. Despite the subtle nature of that tool, significant effects have been discovered at the phenotypic and molecular levels. For instance, deuterium depletion from the normal level of 135-150 ppm to 90 ppm reversed the Mn-induced decrease in *C. elegans* life span through restoring the normality of the DAF-16 pathway ^16^. In another study, administration in the diet of deuterated polyunsaturated fatty acids reduced oxidative stress and extended the lifespan of *C. elegans* ^17^.

The latter result was in agreement with the adage stating that “[isotopically] heavy eaters live longer” ^18, 19^. The origin of this idea can be traced back to James E. Kendall, head of the Chemistry department of the Edinburgh university. This distinguished scientist suggested shortly after the discovery of deuterium in early 1930s that, since deuterium atom is twice as heavy as hydrogen atom, drinking heavy water should slow down all processes, including ageing, and prolong human life span ^20^. On the other hand, depletion of heavy isotopes, e.g. deuterium in ambient water, has been shown to reduce the growth rate of fast-propagating human cells via an increased oxidative stress in mitochondria ^21^. Furthermore, simultaneous depletion of the heavy isotopes of hydrogen, carbon, nitrogen and oxygen in the growth media led to a higher growth rate of bacterium *Escherichia coli* (*E*.*coli*) and 2-3 times faster reactions of the recombinantly expressed enzymes ^22^. Here we investigate the role of heavy isotope depletion on the phenotype and longevity of the nematode *C*.*elegans* feeding on *E. coli* living in isotopically depleted minimal media. Similar to the enzymes expressed in such bacteria ^22^, we will call *C*.*elegans* growing in such conditions ultralight, as their heavy isotopes are significantly depleted compared to the normal (light) isotopic composition.

## RESULTS AND DISCUSSION

### Ultralight *C. elegans* grows faster

The growth of *C. elegans* was assessed via the parameters of the optical density (OD) curves, such as maximum density and maximum growth rate (Fig. 1a). Both parameters indicated that Depleted media provided a more favorable environment for worm growth than Normal media (Fig. 1b). This result was not surprising given our recent finding of the significantly accelerated growth of *E. coli* in the isotopically depleted media ^22^.

**Fig. 1.**
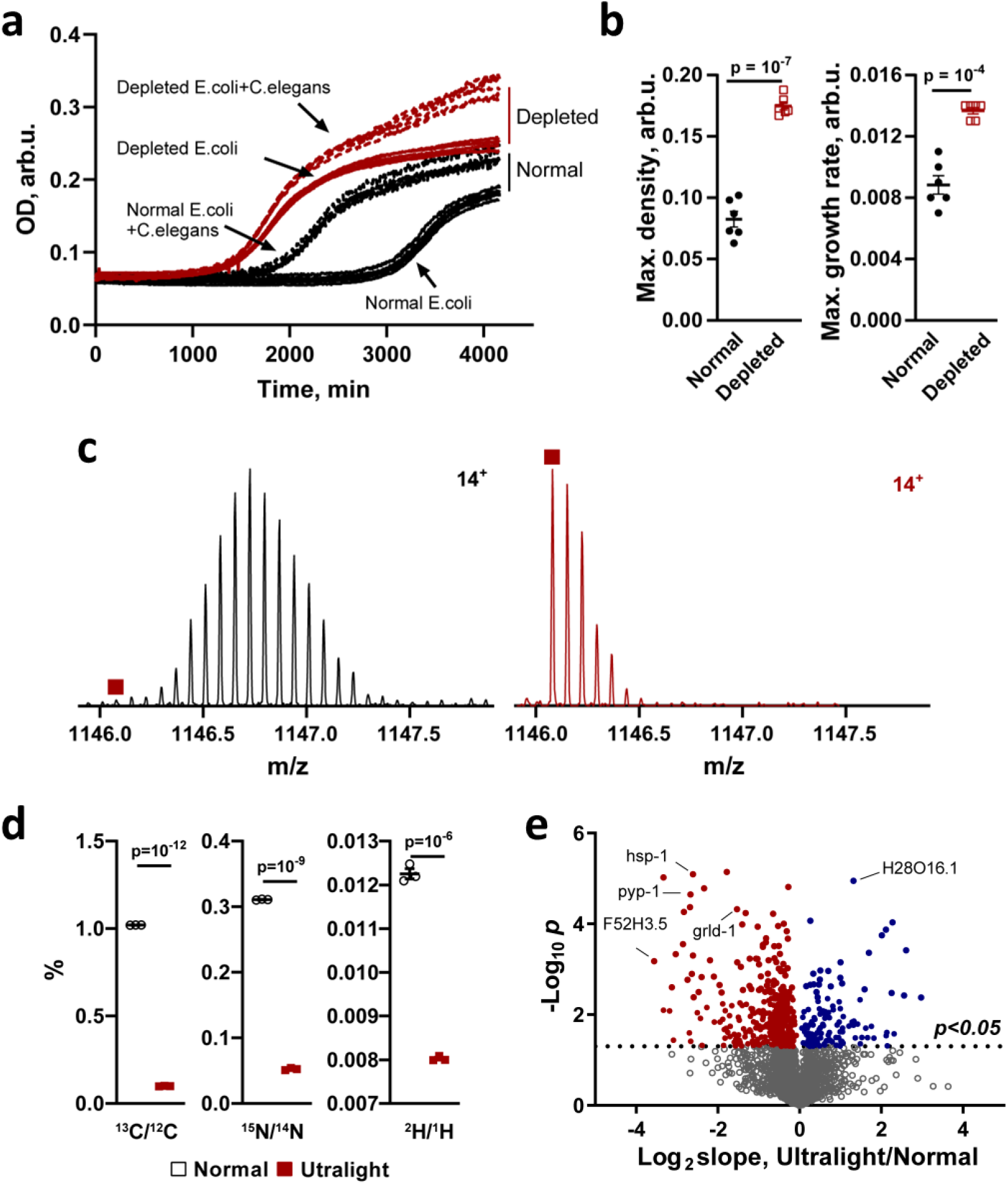
Growth of isotopically normal *C. elegans* grown in Normal media and ultralight *C. elegans* grown in Depleted media and properties of their proteins. **a** The growth curves of *C. elegans* (n=6). **b** Maximum density (left) and maximum growth rate (right) of *C. elegans* (n=6). **c** Mass spectrum of the *C. elegans* protein Galectin (MW 16.1 kDa) grown in the Normal (left) and Depleted (right) media^23^. The monoisotopic mass position is marked by 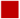 . **d** Isotopic ratio analysis by FT IsoR MS of ^13^C/^12^C (left), ^15^N/^14^N (middle), and H^2^/H^1^ (right) for proteins of *C. elegans* (n=3). **e** Volcano plot of slopes (n=3, median) of the *C. elegans* protein melting curves. The horizontal line indicates the p value of 0.05 in two-tailed unpaired t-test. The ultralight *C. elegans* proteins with steeper slopes have negative Log_2_ values (red dots), while those with more shallow slopes have positive values (blue dots).

### Ultralight *C. elegans* proteins accumulate heavy isotopes

The LC-MS analysis of the worm lysate revealed that, in most proteins with a molecular weight (MW) <20 kDa, the monoisotopic mass dominated in the isotopic distribution of molecular ions (Fig. 1c) ^23^. Fourier Transform Isotopic Ratio Mass spectrometry (FT IsoR MS) ^24^ of the peptide digest obtained from *C. elegans* lysate confirmed profound depletion of ^13^C and ^15^N, as well as a moderate depletion of deuterium in worm proteins (Fig. 1d) compared to normal isotopic composition. However, the depletion of heavy isotopes in worms was much less pronounced than in *E. coli*, their diet. The comparison of the isotope ratio data for the abundant leucine/isoleucine residues (not distinguished due to their isomeric nature) showed that the ^13^C/^12^C ratio increased in *C. elegans* compared to *E. coli* by 124% (p<0.0001), while the ^15^N/^14^N ratio went up by 32% (p<0.002). Both these results are qualitatively consistent with the well-known effect of heavy isotope enrichment along the food chain ^25^. However, the enrichment magnitude is staggering, as normally the enrichment with one trophic level is of the order of 1% for both ^13^C/^12^C and ^15^N/^14^N ^26^. This adds additional arguments to questioning in the isotopic context of the old paradigm “you are what you eat” ^27, 28^.

Curiously, for proline residue the results were different. While the ^13^C/^12^C ratio increased by 61±1.7% (p<10^-6^), the ^15^N/^14^N ratio decreased by 7±1.5% (p<0.03) (Supplementary Fig.1). Although the borderline statistical significance of the latter figure may make it later a subject of revision, these results underline the necessity to study the effect of heavy isotope enrichment with trophic level for each amino acid individually. Note that in the case of ultralight organisms, the traditionally used measure of isotope enrichment relative to the standard expressed in permil (‰) is inapplicable because of the great difference of the analyzed isotope compositions with the established standards.

### Ultralight *C. elegans* proteins show altered thermal stability

Using thermal proteome profiling (TPP), we have previously found that, in *E. coli*, more ultralight proteins increased their thermal stability/solubility compared to the isotopically natural counterparts ^22^. Also, the decrease of stability/solubility with temperature was steeper for ultralight proteins, consistent with their lower vibrational and conformational entropy ^22^. Similar results were observed for *C. elegans* proteins (Fig. 1e, Supplementary Fig. 2a-d, Supplementary Table 1). In analysis of Gene Ontology (GO) processes, proteins with significant differences in both melting temperature (Tm) and melting curve slope (Supplementary Fig. 2e, Supplementary Table 2) were enriched in the pathways determining the adult worm lifespan (Supplementary Fig. 2f). This finding led us to hypothesize that in ultralight *C. elegans* the programmed aging process may be modulated compared to isotopically normal worms.

### Ultralight *C. elegans* develop faster and live shorter lives

The developmental study was conducted by measuring each stage component of the *C. elegans* population every 24 h. Phenotypically, a significant difference emerged as early as at 48 h, with ultralight worms reaching the young adult stage significantly earlier (Fig. 2a).

**Fig. 2.**
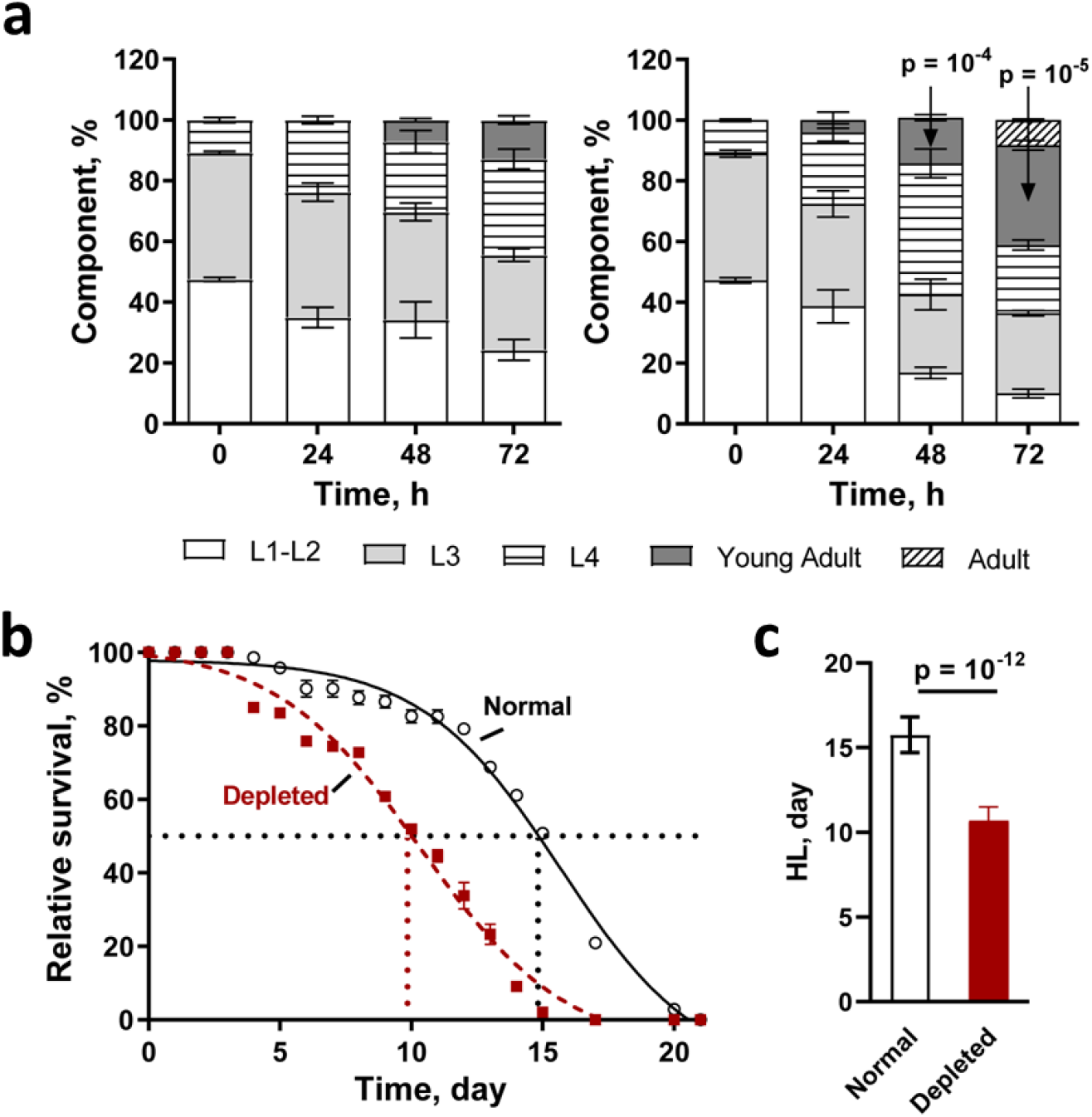
Development and aging of *C. elegans* grown in Normal and Depleted media. **a** Development components of *C. elegans* grown in Normal (left) and Depleted (right) media. **b** The survival curves of *C. elegans* grown in Normal and Depleted media (n=12). **c** Comparison of average lifespans of *C. elegans* from **b** (n=12).

To determine the lifespan, 5-fluoro-2′-deoxyuridine (FUdR) was employed that inhibits DNA synthesis, causing death of larvae and preventing progeny development, not affecting adults. The survival population of *C. elegans* was counted every 24 h. Ultralight worms exhibited a 5.1 ± 0.8 days shorter half-lifespan (HL) compared to normal worms (p <<10^-3^, Fig. 2b-c). These findings supported the hypothesis that the programmed aging and death processes are modulated by isotopic composition of the environment.

### Establishing proteome-based pseudo-time scale of *C. elegans* aging

To get insight into the molecular mechanism of the ultralight *C. elegans* programmed aging and death, we assessed the age-related changes in protein abundance and solubility using the PISA-Express proteomics analysis ^29^. For that, FUdR-treated worms in three biological replicates were harvested on day 0 and day 10 (for both conditions) as well as day 15 (for normal worms only), with the first time point representing young age and the last two - adulthood (Fig. 3a). For each replicate, cell lysates were divided into two parts: one for protein expression analysis and the other for protein solubility analysis. In the latter analysis, 12 aliquots were subjected to treatment at a different temperature, ranging from 48 °C to 59 °C, with a 1 °C interval before aliquot pooling, ultracentrifugation, and collection of soluble proteins for digestion. The resulting digests were analyzed by liquid chromatography-tandem mass spectrometry (LC-MS/MS), and protein abundances (Supplementary Table 3) as well as thermally induced changes in protein solubility (Supplementary Table 4) were assessed.

**Fig. 3.**
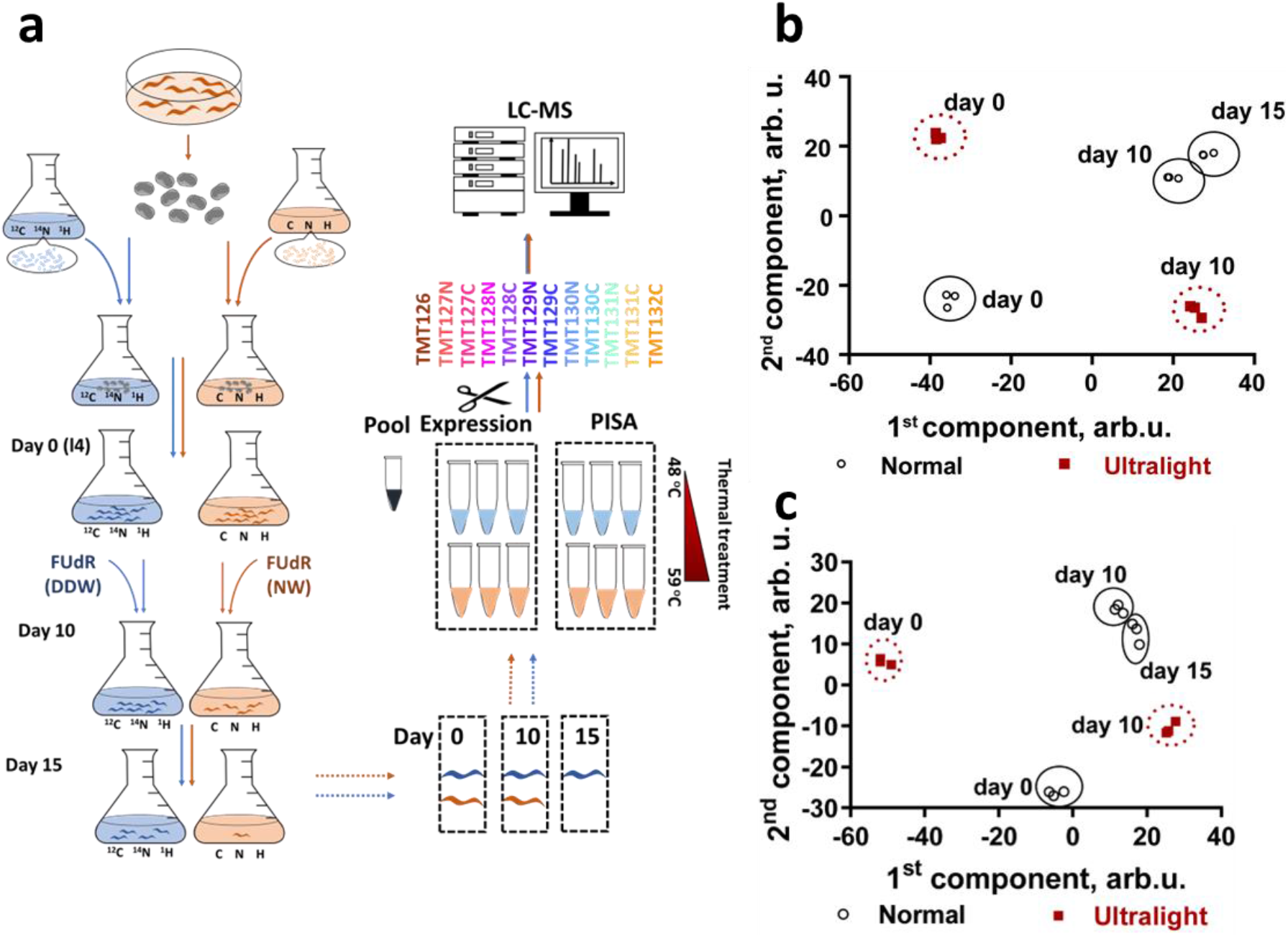
Establishing a pseudo-time scale for aging of *C. elegans* grown in Normal and Depleted media. **a** Workflow of PISA-Express analysis of worm proteins. PCA plot of protein abundance (**b**) and solubility (**c**) changes during aging for normal and ultralight *C. elegans* (n=3).

Principal Component Analysis (PCA) of protein expression and, separately, protein solubility was performed, with the results mapped on a 2D plot defined by the first two principal components (Fig. 3b, c). As the first components in both analysis domains correlated with age, they were interpreted to be pseudo-time scales for worm ageing. The expression-based pseudo-time scale (Fig 3b) showed that at day 0, both normal and ultralight worms were, not surprisingly, of the same “age”. However, by day 10 ultralight *C. elegans* were “older” than 10-days old normal worms, although somewhat “younger” than their 15-day old normal counterparts. This result was unexpected, given that in terms of survival rate the 10-days old ultralight worms are equivalent to the 15-days old normal *C. elegans* (Fig. 2b).

The protein solubility analysis provided even more surprising results (Fig. 3c). According to the pseudo-time protein solubility scale, the ultralight C. elegans are much “younger” than their isotopically normal counterparts at day 0, and much “older” at day 10 than even the 15-days old normal worms.

It should be noted that this is not the first time the solubility analysis provides greater separation between different biological states than the expression analysis; a similar effect was observed in studying stem cell differentiation by PISA-Express ^29^. This result underlines the importance of studying the biological processes in both analytical domains.

### Protein solubility difference during aging

Given that protein solubilities seem to be more informative than protein abundances for delineating the ageing trajectory, protein solubilities were compared for ultralight versus normal *C. elegans* for day 0 (Fig. 4a, b) and day 10 (Fig. 4c, d). The top 50 proteins with largest solubility differences for day 0 were most significantly enriched in translation (Fig. 4b). Additionally, mitochondrial electron transport was identified as one of the significant biological processes, with many of the proteins located in the mitochondria. Thus, mitochondrial electron transport, along with ROS generation and uncoupling, may contribute to the differences observed between the young normal and ultralight *C. elegans* (Supplementary Fig. 3a). For day 10, proton transmembrane transport, closely associated with ROS generation, was the most significantly enriched process for the top 50 proteins with largest solubility differences. Again, many of these proteins were mapped to the mitochondria (Supplementary Fig. 3b).

**Fig. 4.**
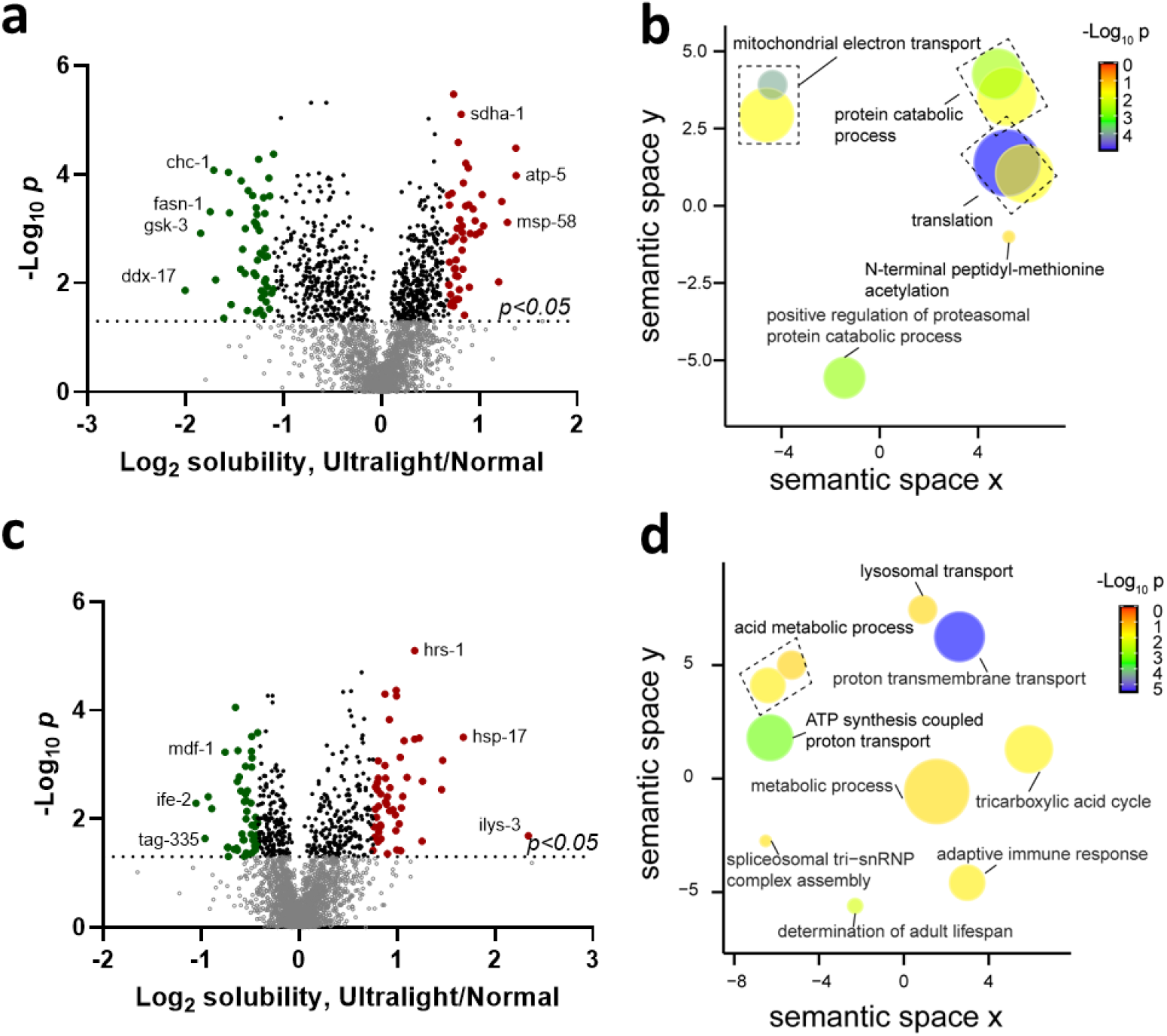
Protein solubility differences for *C. elegans* grown in Normal and Depleted media. **a** Volcano plot, and **b** GO enrichment in biological process at day 0. **c, d** Same as **a** and **d**, respectively, for day 10. The horizontal line indicates the p value of 0.05 in two-tailed unpaired t-test. GO term enrichment in biological process (**b, d**) performed using DAVID and plotted using REVIGO. The size of the bubbles is indicative of the number of proteins annotated with that GO term; bubbles are color coded according to significance.

### Oxidation-reduction processes are preferentially affected during aging

The above findings pointed to the importance of mitochondrial processes and ROS generation-related activities in understanding the variation in *C. elegans* lifespan under different isotopic conditions. Indeed, increased cellular ROS levels are known to accelerate aging processes in cells ^30^. Also, deuterium depletion in ambient water increased the ROS generation in human cells ^21^. Thus we hypothesized that redox processes are implicated in the shortening of the worm lifespan due to heavy isotope depletion. In order to test this hypothesis, we measured ROS production in *C. elegans* on a daily basis (Fig. 5). Ultralight worms exhibited higher ROS production starting from day 0, with the degree of overproduction increasing with their age. The relative ROS production reached its first peak corresponding to ≈200% of ROS levels in normal worms on day 4; this was the day preceding the onset of mortality in ultralight worms. On day 13, when the difference in the survival of *C. elegans* was most pronounced, the relative ROS production in ultralight worms peaked at the maximum value of ≈400% compared to normal worms. This result verified the hypothesis that enhanced ROS levels are responsible for the life shortening effects in ultralight *C. elegans*.

**Fig. 5.**
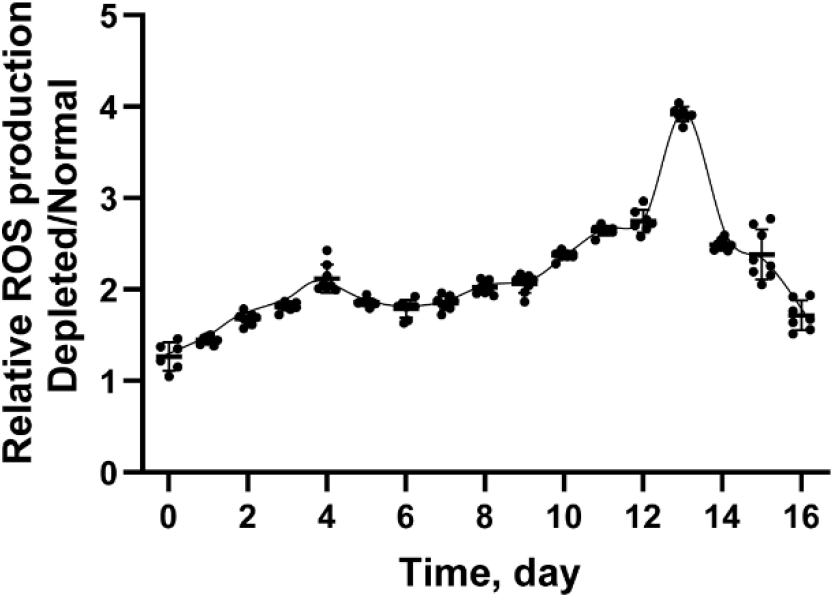
ROS overproduction in Ultralight worms. Relative ROS production in *C. elegans* grown in Depleted media compared with Normal media during aging (n=8).

## CONCLUSION

Here we showed for the first time that heavy isotope depletion of the environment, particularly food, may greatly affect not only the growth rate, but also longevity of animals. Depletion of heavy isotopes in the current work puts it aside from the earlier studies, conducted mostly in 1960s and 1970s, on a >50% enrichment with heavy isotopes ^13^C ^31^ and ^18^O ^15^ as well as more recent studies on ^15^N enrichment ^32^. While depletion in our experiments affected less than 1% of all atoms, the effects observed were much greater than this ratio might suggest in a naïve linear approximation. However, in the wake of our recent finding that ultralight enzymes can exhibit 2-4 times higher activity compared to isotopically normal counterparts ^22^, our current results do not look completely unexpected. The very significant heavy isotope enrichment in some amino acids of *C. elegans* proteins compared to severely isotopically depleted *E. coli* that the worms use as food warrants further thorough investigation. We hope that this work provides solid justification for the future extensive studies of the active role stable isotopes play in life sciences.

## ACKNOWLEDGEMENTS

Swoboda lab at Karolinska Institutet, Stockholm is acknowledged for providing *C. elegans*. This work was funded by the Swedish Research Council, grant 2021-05223 to RAZ. RAZ also acknowledges support from The Ministry of Science and Higher Education of the Russian Federation (agreement no. 075-15-2020-899). A portion of this work was conducted at the National High Magnetic Field Laboratory ICR User Facility, which is funded by the National Science Foundation Division of Chemistry through DMR-1644779 and the State of Florida.

## Methods

### Media Preparation

M9 minimal media were prepared as follows. Na_2_HPO_4._2H_2_O, KH_2_PO_4_, NaCl, NH_4_SO_4_, MgSO_4_, CaCl_2_ and glucose were obtained from Sigma-Aldrich. ^13^C-depleted glucose was purchased from Cambridge Isotope Laboratories and ^15^N-depleted (NH_4_)_2_SO_4_ was from Merck. Deuterium Depleted Water (DDW) containing ≤5 ppm D and 410 ppm ^18^O was obtained from MTC Iceberg Ltd (Moscow, Russia). M9 stock salt solutions (M9 5×SS) were prepared by dissolving 42.5 g Na_2_HPO_4._2H_2_O, 15.0 g of KH_2_PO_4_ and 2.5 g of NaCl in 1000 mL of either normal milli-Q water (for Normal M9 media) or DDW (for Depleted M9 media). Both solutions were autoclaved before proceeding to M9 media preparation. M9 media were prepared by mixing 800 mL of either milli-Q water (for Normal M9 media) or DDW (for Depleted M9 media), 200 mL of the corresponding M9 5xSS, 2.0 mL of 1M MgSO_4_, 0.1 mL of 1M CaCl_2_, 5 g of glucose and 1 g of (NH_4_)_2_SO_4_ (isotopically normal for Normal M9 media and ^13^C-depleted glucose and ^15^N-depleted (NH_4_)_2_SO_4_ for Depleted M9 media). These media were filtered using 0.2 μm polyether sulfone (PES) filters (VWR) before use.

#### *C. elegans* growth

For each set of experiments, *C. elegans* N2 strain with mixed worm population were grown on plates containing Nematode Growth Medium (NGM) with *E. coli* OP50. The worms were collected and washed with M9 media and transferred to Normal or Depleted M9 media. *E. coli* BL21 grown in Normal M9 media were further diluted (1:500) into Normal or Depleted M9 media containing *C. elegans* and put into honeycomb 100-well plates (BioScreen, Finland) in 6 replicates. Both bacterial and worm growth was monitored by measuring optical density on a BioScreen C instrument (BioScreen, Finland) at 20 °C. The number of worms at each stage was recorded every 24 h.

#### *C. elegans* lifespan

*C. elegans* N2 strain were grown on plates containing NGM with *E. coli* OP50 and collected. Then the worms were washed with the same type of M9 media (Normal or Depleted) in which *E. coli* were growing and lysed using freshly prepared sodium hypochlorite solution by vertexing until worm bodies are dissolved and only eggs are obvious under the dissecting microscope. The egg preparation was spun down for 2 min at 300×g, washed three times with sterile water and resuspended in Normal or Depleted M9 media. After incubation for overnight, the number of L1 larvae were counted under microscope and the final volume was adjusted so that worm concentration in the suspension was approximately 100 worms/mL. Feeding bacteria grown in Normal or Depleted M9 media were added in the suspension to a final concentration of 0.5 mg/mL. Afterwards, 100 μL of L1 larvae suspension containing bacteria was added in each well of a 96-well plate and incubated at 20 °C. When all worms transferred to L4 larvae, FUdR (F0503, Sigma) dissolved in Normal or DDW was added into each well to a final concentration of 120 μM. The plate was incubated at 20 °C and the number of living and dead worms in each condition was recorded under the dissecting microscope every day.

### Measurement of Cellular ROS Concentration

The worms grown in Normal and Depleted M9 media were collected every day and washed by M9 media and DCF-DA in DMSO was added into the suspension containing *C. elegans* to the final concentration of 20 μM. After 30 min incubation at 20 °C in darkness, the fluorescence intensity was measured by Infinite® M200 PRO (TECAN, Männedorf, Switzerland). The excitation and emission wavelengths were 485 nm and 535 nm, respectively. In parallel, the relative worm survival was observed under microscope. The fluorescence intensity in each well was normalized to the average number of worms on the same day.

### TPP sample preparation

Mixed *C. elegans* populations were grown in Normal and Depleted M9 media. After 72 h of growth, the worms in each flask were collected, washed, resuspended in PBS with protease inhibitor (5892791001, Sigma) and lysed by probe sonication. The protein solution was collected after centrifuge and divided into 8 aliquots. These aliquots were incubated for 3 min at either 37, 43, 49, 55, 61, 67, 73 or 79 °C. After that the samples were kept at room temperature (RT) for 5 min to cool down and ultra-centrifuged at 35,000 rpm/min at 4 °C for 30 min. Afterwards, the supernatant was collected, reduced with 10 mM DTT (10708984001, Sigma) and alkylated with 25 mM IAA (I1149, Sigma). The samples were precipitated using cold acetone at -20 °C overnight, digested by Lys C (125–05061, Wako Chemicals GmbH) at a 1:75 enzyme to protein ratio for 6 h at 30 °C and then by trypsin (V5111, Promega) (1:50 enzyme to protein ratio) overnight. After labeling using 16 TMTpro reagents (A44520, Thermo Fisher Scientific) according to manufacturer’s instructions, multiplexing and desalting with C18 Sep-Pak columns (WAT054960, Waters), the peptides samples were fractionated using a Dionex Ultimate 3000 UPLC system (Thermo Fisher Scientific). Every 8th fractions were combined to a single pool and 12 such pools for each TMT-multiplexed sample set were analyzed by nanoLC-MS/MS as described before ^22^.

### PISA-EXPRESS sample preparation

The worms grown in Normal and Depleted M9 media were collected at days 0, 10 and 15, washed and resuspended in PBS with protease inhibitor and lysed by probe sonication. The protein solution was collected after centrifugation and divided into 12 aliquots. Each aliquot was incubated for 3 min at a temperature in the range 48-59 °C, with a 1 °C interval. After that the samples were kept at RT for 5 min to cool down, the protein samples in the same replicate were combined and ultra-centrifuged at 35,000 rpm/min at 4 °C for 30 min. Then the samples were digested, labelled by 16 TMTpro reagents, fractionated and analyzed by nanoLC-MS/MS as above.

### NanoLC-MS/MS analysis

NanoLC-MS/MS analyses were performed on an Orbitrap Fusion Lumos mass spectrometer (Thermo Fisher Scientific). The instrument was equipped with an EASY ElectroSpray source and connected online to an Ultimate 3000 nanoflow UPLC system. The samples were pre-concentrated and desalted online using a PepMap C18 nano-trap column (length - 2 cm; inner diameter - 75 μm; particle size - 3 μm; pore size - 100 Å; Thermo Fisher Scientific) with a flow rate of 3 μL/min for 5 min. Peptide separation was performed on an EASY-Spray C18 reversed-phase nano-LC column (Acclaim PepMap RSLC; length - 50 cm; inner diameter - 2 μm; particle size - 2 μm; pore size – 100 Å; Thermo Scientific) at 55 °C and a flow rate of 300 nL/min. Peptides were separated using a binary solvent system consisting of 0.1% (v/v) FA, 2% (v/v) ACN (solvent A) and 98% ACN (v/v), 0.1% (v/v) FA (solvent B). They were eluted with a gradient of 3–26% B in 97 min, and 26– 95% B in 9 min. Subsequently, the analytical column was washed with 95% B for 5 min before re-equilibration with 3% B. The mass spectrometer was operated in a data-dependent acquisition mode. A survey mass spectrum (from m/z 375 to 1500) was acquired in the Orbitrap analyzer at a nominal resolution of 120,000. The automatic gain control (AGC) target for was set as 100% standard, with the maximum injection time of 50 ms. The most abundant ions in charge states 2^+^ to 7^+^ were isolated in a 3 s cycle, fragmented using HCD MS/MS with 33% normalized collision energy (NCE), and detected in the Orbitrap analyzer at a nominal mass resolution of 50,000. The AGC target for MS/MS was set as 250% standard with a maximum injection time of 100 ms, whereas dynamic exclusion was set to 45 s with a 10-ppm mass accuracy window.

### Bottom-up proteomics data analysis

The LC-MS/MS raw files were processed by an in-house modified version of MaxQuant software (version 1.6.2.3) recognizing TMTpro as an isobaric mass tag using the “Specific Trypsin/P, Lyc/P” digestion mode with maximum two missed cleavages as described before ^22^. The MS/MS spectra were searched against the Uniprot *Caenorhabditis elegans* Bristol N2 database (UP000001940, containing 26,695 entries, last modified on August 3, 2020).

### TPP data analysis

For curve fitting of TPP experiments an in house developed R package was used, publicly available on GitHub (https://github.com/RZlab/SIESTA). In short, proteins with fewer than two peptides, contaminants, proteins with reversed sequences, or proteins with missing abundance values in any of the three replicates were removed from further analysis. The protein melting curves were fitted with the equation and the middle point provides the melting temperature Tm:

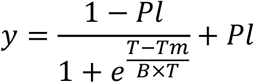

where the term B is responsible for the slope steepness (lower values mean steeper curve) and Pl is the high-temperature plateau of the melting curve ^33, 34^. For statistical analysis of slopes and Tm, two-sided t-tests (with equal or unequal variance depending on F-test) were applied.

### PISA-EXPRESS data analysis

After removal of contaminates, reverse, proteins identified only by site and proteins with less than two peptides, each TMT channel was normalized by its total abundance. For solubility analysis, the value for each protein was first normalized by the mean expression value for all replicates in order to remove any abundance bias. Statistical analysis was done by using a two-tailed Student’s t-test with equal or unequal variance depending on F-test.

### FT IsoR MS analysis

The cell lysate was digested as above, and after desalting the peptides were analyzed by nanoLC-MS/MS. Mass spectra were acquired on an Orbitrap Fusion Lumos Mass in the data-independent acquisition mode with peptides in charge state ≥2^+^ selected for MS/MS in an isolation window 1000 m/z units wide centered at m/z 800. MS/MS was performed with HCD energy set at 50 NCE. The detection range was from m/z 50 to 200, with the nominal mass resolving power of 60,000. Data processing was done using the in-house developed Github package, PAIR-MS, analyzing the fine isotopic structure of the immonium ions of amino acids Pro, Val and Leu/Ile, as described earlier ^25^.

## Data availability

Excel files containing the analyzed data are provided in Supplementary Materials. The mass spectrometry proteomics data have been deposited to the ProteomeXchange Consortium (http://proteomecentral.proteomexchange.org) via the PRIDE partner repository with the dataset identifier PXD049006.

## Code availability

The curve fitting R package is available in GitHub (https://github.com/RZlab/SIESTA) and the package for FT IsoR MS analysis is available in GitHub (https://github.com/RZlab/isoms).

