## Supporting Information for "Do light eaters live shorter lives? The case of ultralight *Caenorhabditis elegans*"

**Do lighter eaters live shorter lives?**

**The case of ultralight *Caenorhabditis elegans***

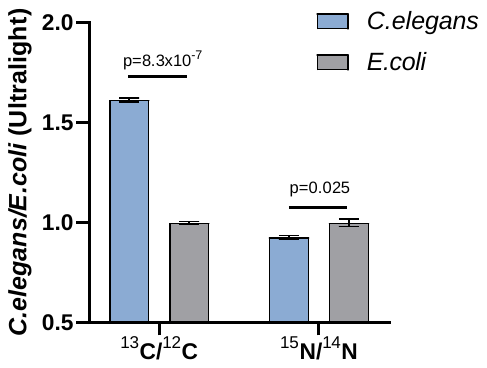

**Supplementary Fig. 1 Ratio of ^13^C/^12^C and ^15^N/^14^N in proline residue of ultralight proteins for *C. elegans* compared with *E. coli* (n=3).**

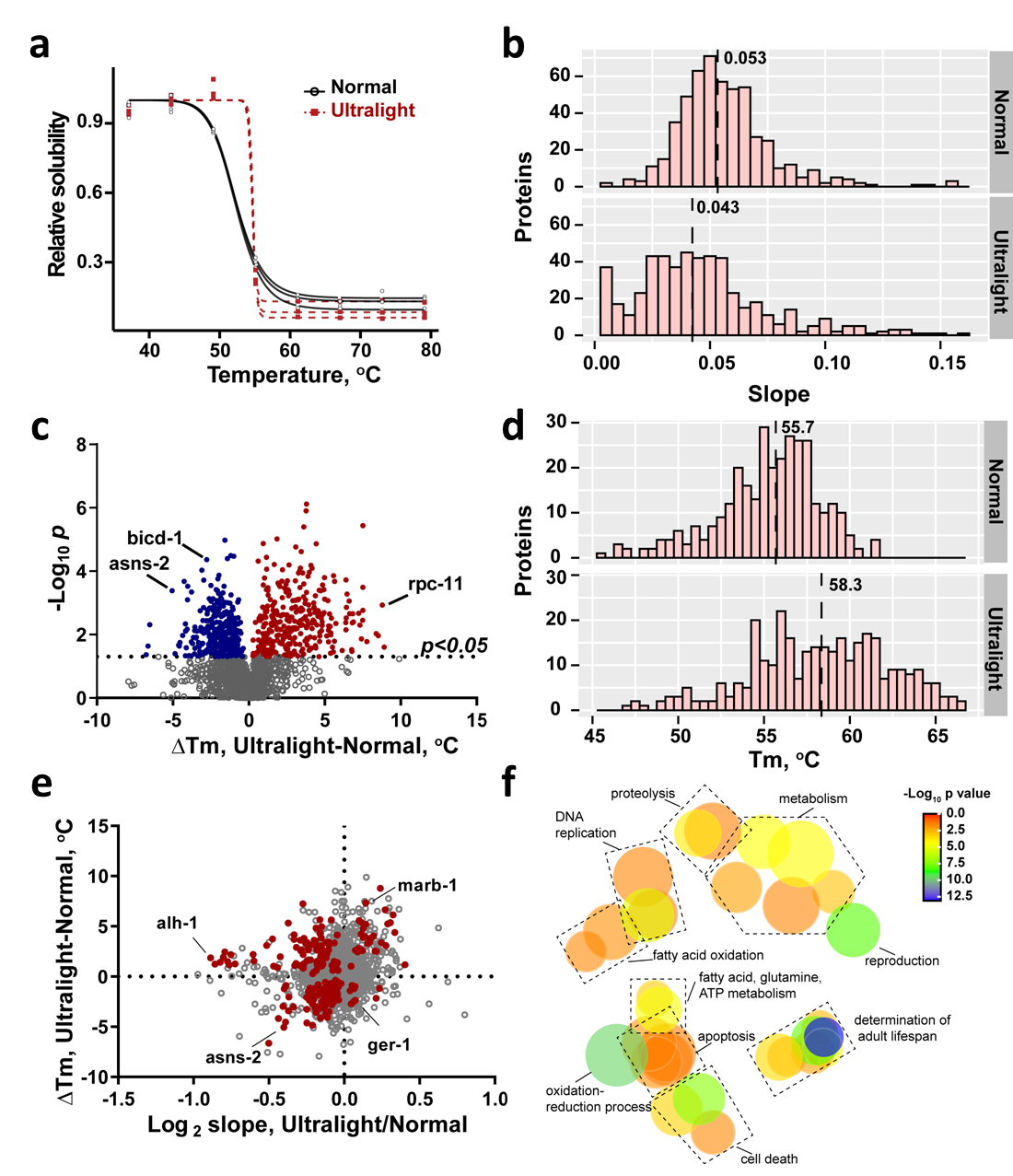

**Supplementary Fig. 2 Comparison of thermal stability parameters of proteins from isotopically normal *C. elegans* grown in Normal media and ultralight worms grown in Depleted media. a** The melting curves of alh-1 protein (n=3). **b** Histogram of significantly different slopes of protein melting curves. The lines indicate the median slopes values. **c** Volcano plot of differences in protein melting temperature Tm (n=3, medians). The horizontal line indicates the p value of 0.05 in two-tailed unpaired t-test. The ultralight proteins more stable than normal counterparts have positive ∆Tm values (red dots), while less stable proteins have negative ∆Tm values (blue dots). **d** Histogram of proteins with significantly different melting temperatures Tm. The lines indicate the median Tm values. **e** Correlation between melting temperatures and melting curve slopes of *C. elegans* proteins (n=3, medians). The proteins with significant differences in both Tm and slope are labeled as red dots. **f** GO term enrichment (biological process) of proteins with significant difference in both Tm and melting curve slopes performed using DAVID and plotted using REVIGO. The size of the bubbles is indicative of the number of proteins annotated with that GO term; bubbles are color-coded according to significance of the term.

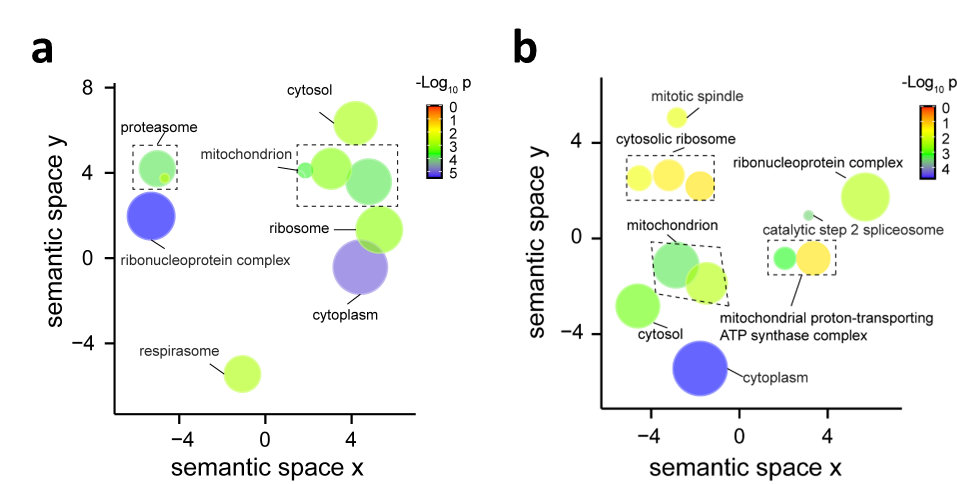

**Supplementary Fig. 3** Go enrichment (cellular component) of top 50 proteins with largest solubility differences in ultralight vs normal *C. elegans* at **a** day 0, and **b** day 10. GO term enrichment in molecular component performed using DAVID and plotted using REVIGO. The size of the bubbles is indicative of the number of proteins annotated with that GO term; bubbles are color coded according to significance.
